# A novel *in silico* scaffold-hopping method for drug repositioning in rare and intractable diseases

**DOI:** 10.1101/2023.07.03.547598

**Authors:** Mao Tanabe, Ryuichi Sakate, Jun Nakabayashi, Kyosuke Tsumura, Shino Ohira, Kaoru Iwato, Tomonori Kimura

**Affiliations:** Laboratory of Rare Disease Information and Resource library, Center for Intractable Diseases and ImmunoGenomics Research, National Institutes of Biomedical Innovation, Health and Nutrition (NIBIOHN), Ibaraki-city Oosaka, Japan; Analysis Technology Center, FUJIFILM Corporation, 210 Nakanuma, Minamiashigara-city Kanagawa, Japan; Reverse Translational Research Project, National Institutes of Biomedical Innovation, Health and Nutrition (NIBIOHN), Ibaraki-city Oosaka, Japan; KAGAMI Project, National Institutes of Biomedical Innovation, Health and Nutrition (NIBIOHN), Ibaraki-city Osaka, Japan

## Abstract

In the field of rare and intractable diseases, new drug development is difficult and drug repositioning (DR) is a key method to improve this situation. In this study, we present a new method for finding DR candidates utilizing virtual screening, which integrates amino acid interaction mapping into scaffold-hopping (AI-AAM). At first, we used a spleen associated tyrosine kinase (SYK) inhibitor as a reference to evaluate the technique, and succeeded in scaffold-hopping maintaining the pharmacological activity. Then we applied this method to five drugs and obtained 144 compounds with diverse structures. Among these, 31 compounds were known to target the same proteins as their reference compounds and 113 compounds were known to target different proteins. We found that AI-AAM dominantly selected functionally similar compounds; thus, these selected compounds may represent improved alternatives to their reference compounds. Moreover, the latter compounds were presumed to bind to the targets of their references as well. This new “compound-target” information provided DR candidates that could be utilized for future drug development.

## Introduction

Approximately 7000 rare and intractable diseases (RIDs) have been defined to date, affecting an estimated 300 million people worldwide^1^. These diseases largely reduce patients’ quality of life throughout their lives. Though the unmet medical needs are very high in this field, new drug development for RIDs is difficult. The reason is that the number of affected patients is too small for pharmaceutical companies to invest in targeting these diseases^2, 3^ and the mechanism of onset of many RIDs still remains to be elucidated. Therefore, attention is focused on drug-repositioning (DR) methods, finding a candidate drug previously developed for other diseases^4, 5, 6^. When relatively little information is available for the disease, phenotypic screening and target-based method are selected from existing DR methods^5^. We considered that a drug which was rather effective in a RID and known to target some protein, even when its action mechanism was not thoroughly known, could be replaced by improved alternative drug using target-based method. Target-based methods include *in vitro* and *in vivo* high-throughput screening of drugs and *in silico* (computational) screening of drugs from libraries^7, 8^. Computational methods, virtual screening (VS) techniques, are generally classified into two major categories: structure-based virtual screening (SBVS) and ligand-based virtual screening (LBVS). SBVS encompasses methods that exploit the three-dimensional (3D) structure of the target and molecular docking^9, 10^. LBVS can be employed when the target structure of sufficient quality for docking simulations is not available and some binders for the target binding pocket are already known^11, 12^. LBVS mainly includes methods based on similarity, in which the relationships between compounds in a given library and known binding molecules for the target are examined by similarity measurements using suitable molecular descriptors^13^. To find compounds that are structurally diverse but share some biological activity, scaffold-hopping, which is a LBVS approach, has been widely attempted^14^. However, because only a few active ligands are available to be used as references, the hit compounds found by LBVS lack novelty^15, 16^. As a wide diversity of hit structures is important for improved properties, some hybrid strategies that integrate both SBVS and LBVS techniques have been proposed to overcome the weakness of LBVS^16, 17^. In this study, we developed a new methodology called AI-AAM, which enabled obtaining candidates with a wide variety of structures by using only the ligand-based virtual scaffold-hopping. Our hypothesis was that the interactions between a ligand and the set of amino acids could represent the interaction between a ligand and its target protein. By introducing Amino Acid Mapping (AAM), the descriptor of the interactions of a compound with amino acids, to the scaffold-hopping technique, we aimed to discover the compounds that have preserved interactions with their targets.

In this report, we aimed to examine the possibility of DR using AI-AAM with 6 compounds as reference. An overview of this study is shown in Fig. 1. At first, we selected a SYK inhibitor as reference to evaluate the technique experimentally, focusing on the pharmacological activity of a hit with the different scaffold from the reference. Then we applied this method to other five reference compounds selected from DDrare, a database of Drug Development for Rare Diseases, and examined whether the hit compounds were structurally diverse and target the same proteins as reference compounds. Moreover, on the basis of the target information of hit compounds, we investigated the pharmacological functions of hits or inferred the new compound-target connection. Last, we discuss the possibility of DR using AI-AAM, based on the results.

**Figure 1:**
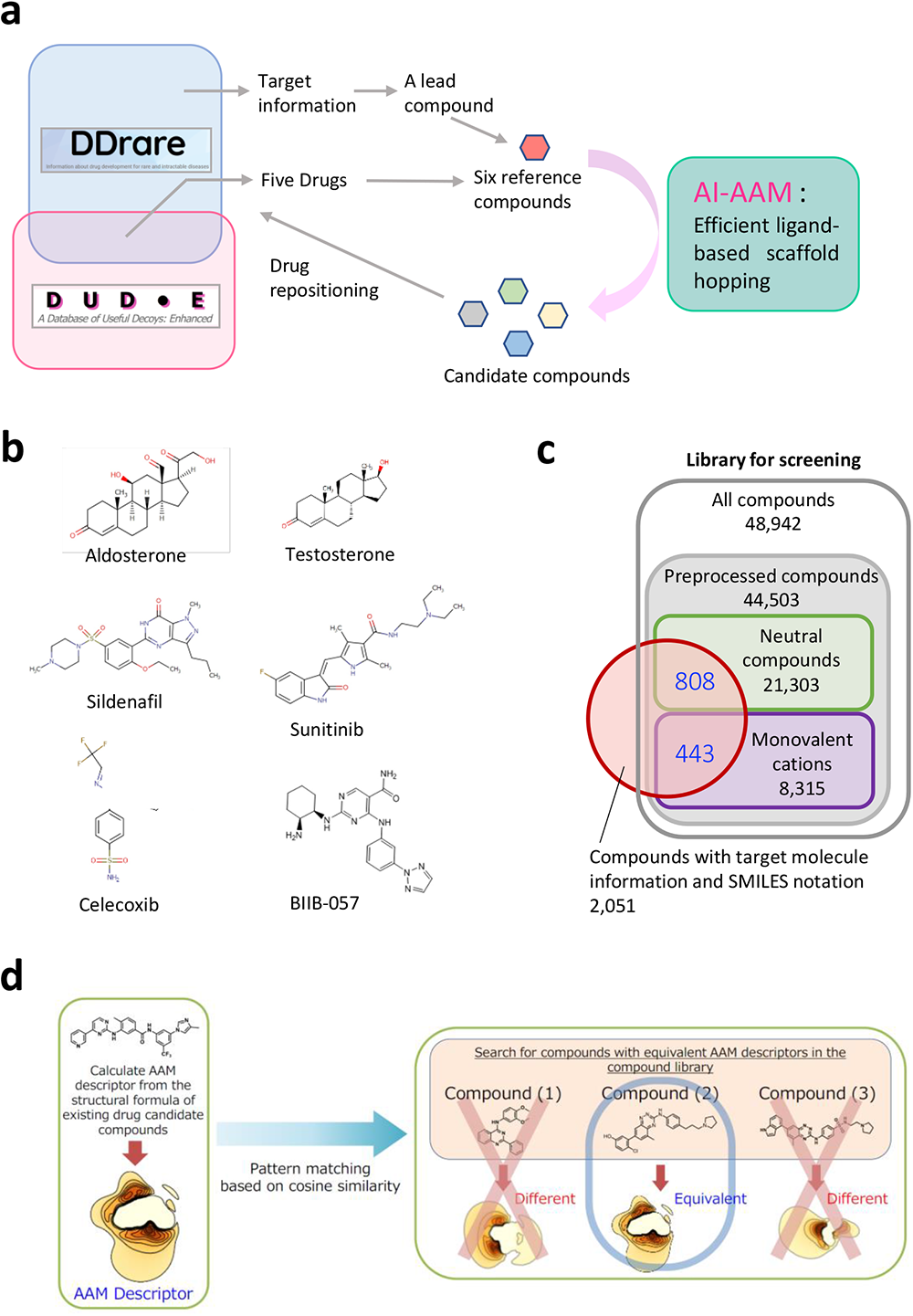
Schematic representation of this study. **a** A lead compound, based on the target information of DDrare, and five drugs contained in both DUD-E and DDrare were selected as reference compounds. Chemical libraries were then explored by scaffold-hopping using AI-AAM. Identified compounds are candidates for drug repositioning for rare and intractable diseases. **b** The structural formulae of six reference compounds. **c** The range of the search for novel hit compounds. The neutral compounds and the monovalent cations including the five reference compounds constitute a part of the Namiki compound library set for repositioning (Namiki Shoji Co., Ltd.), which has 48,942 types in total. The number of the compounds preprocessed successfully was 44,503. Among these compounds, those that are also registered in the DrugBank and written in SMILES notation were selected. The number of these compounds is 1251 that consisted of 808 neutral and 443 monovalent cation compounds. Within the limits of these compounds, the search for novel hits was performed. **d** Schematic representation of virtual screening by AI-AAM.

## Results

### The search for compounds using the AI-AAM technique

AI-AAM was applied to the drugs in the field of RIDs or in clinical trials as reference compounds. Information regarding the drugs was obtained from DDrare, a database of drugs and their target information used in clinical trials of rare diseases (https://ddrare.nibiohn.go.jp/) (Fig. 1a). For the experimental validation of the technique, we selected a known SYK inhibitor candidate BIIB-057 as reference on the basis of target information in DDrare. Then, for the detailed analysis of the characteristics of the screening, we chose 5 compounds (i.e., aldosterone, testosterone, sildenafil, sunitinib and celecoxib) from the compounds registered with both DDrare and Directory of Useful Decoys, Enhanced (DUD-E), as reference (Fig.1a, b). DUD-E is a database of useful decoys designed to help benchmark molecular docking programs^18^. Based on these 6 compounds, a search for DR candidate compounds was performed using AI-AAM (Fig. 1a). In the chemical library, 44,503 compounds were preprocessed successfully and subjected to screening by AI-AAM, among which 1251 compounds (neutral compounds, 808; monovalent cations, 443) had target information in DrugBank (https://go.drugbank.com/) and were used for comparative analysis of target (Fig. 1c). For both the reference and candidate compounds, AAM descriptors, which describe the set of interactions between amino acids and the compound, were calculated and the compounds with similar AAM descriptors were screened from compound libraries and identified as hits (Fig.1d, see “Calculations of AAM descriptors” in Methods for more details). The hits were analyzed and validated in terms of comprehensiveness and specificity.

### Experimental validation for a hit compound identified with AI-AAM

As mentioned above, we selected a known SYK inhibitor candidate BIIB-057 as reference to discover other lead compounds. SYK is a non-receptor tyrosine kinase associated with many RIDs and is considered to be a worthy drug target. With BIIB-057 as the reference, 18 compounds with similar AAM descriptors were identified, and one of them, XC608, which had a scaffold that differed from the reference, was selected to evaluate the levels of inhibitory activity for the target, SYK (Fig. 2a). The IC50 values for BIIB-057 and XC608 were 3.9 nM and 3.3 nM, respectively. These values are close to each other, suggesting that XC608 inhibits SYK activity as effectively as BIIB-057. These results indicate that the compounds predicted to have similar pharmacological activity to their reference compound by AI-AAM have also experimentally obtained IC50 values, which represent the potency of the inhibitor, close to the reference.

**Figure 2:**
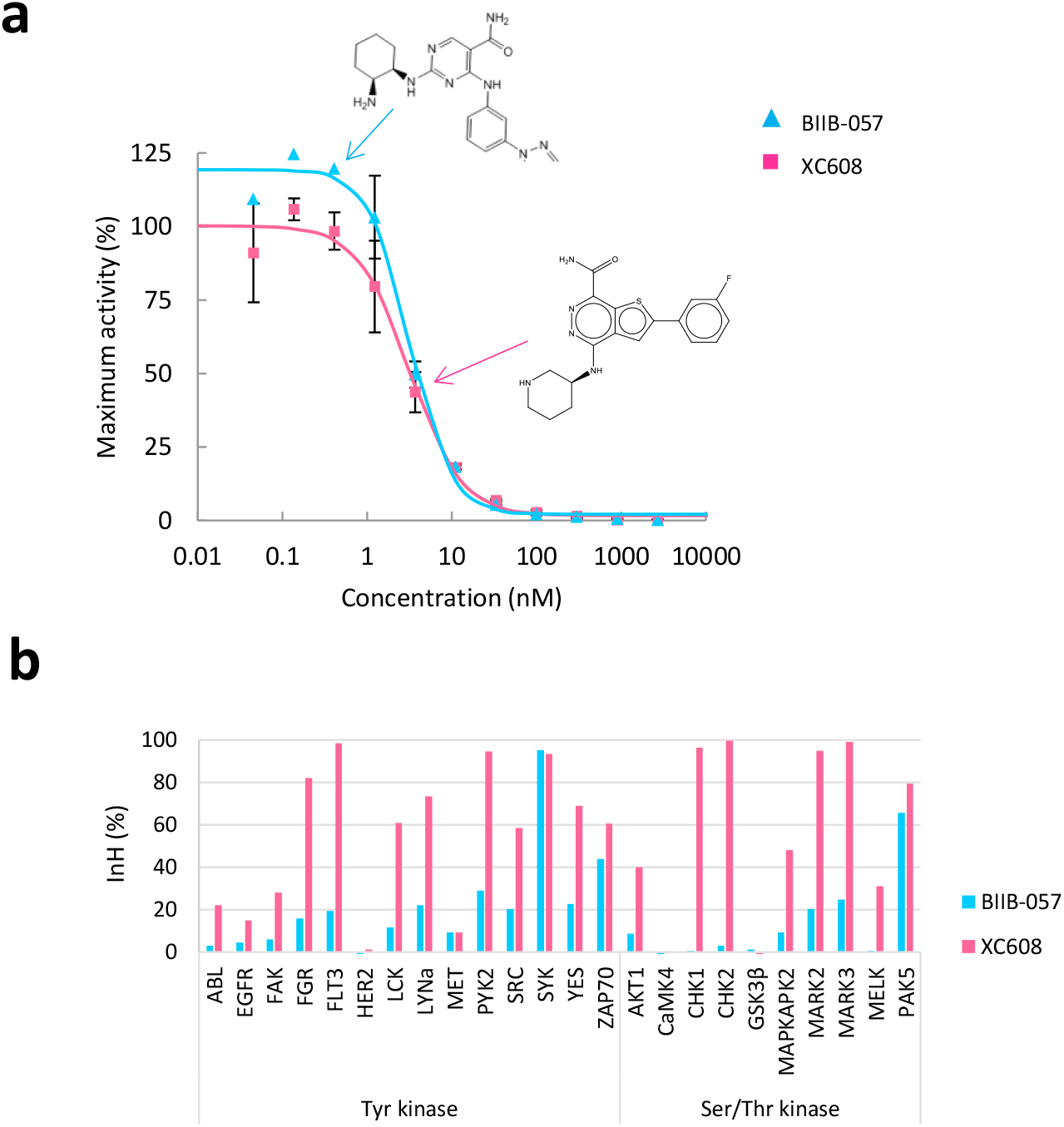
Comparison between BIIB-057, the reference, and XC608, the hit, in inhibitory activity for kinases. **a** SYK kinase activity dose–response curves using BIIB-057 and XC608. The kinase activity IC50 values for BIIB-057and XC608 are 3.9nM and 3.3nM, respectively. **b** Kinase profiling of BIIB-057 and XC608 using mobility shift assay (ATP concentration: Km value of each kinase, Concentration of each compound: 50 nM).

Next, kinase profiling of BIIB-057 and XC608 was performed to examine their selectivity. Among 24 kinases examined, 2 and 14 were inhibited at least 50% by BIIB-057 and XC608, respectively (Fig. 2b). These results suggest that BIIB-057 inhibits SYK and PAK5 selectively, while XC608 inhibits many kinases including SYK with a lower selectivity.

These results showed that the screening setting the threshold value of AAM similarity scores for 0.7 provided the compounds with different structure and selectivity than the reference, maintaining the pharmacological activity.

### Comprehensiveness of screening using AI-AAM

A total of 1275 compounds comprising 25 to 488 new compounds per reference compound, were obtained as hits. When the compounds were limited to those registered in DrugBank, there were 144 compounds identified in total, with 2 to 70 based on each reference (Table 1). The compounds registered in DrugBank have target information, and Table 1 shows the result of aggregation according to whether their known targets are the same as those of the reference compounds.

We estimated how comprehensively AI-AAM identified the compounds whose known targets were the same proteins as the reference compounds. After all compounds that were known to target the same proteins as the references were counted, we examined how many of these compounds were obtained as hits using AI-AAM (Table 1 “Extraction rate”). For aldosterone, testosterone, and sildenafil, the extraction rate was greater than 60%. For sunitinib, the extraction rate was 33.3% when the target was limited to KIT only, whereas it was 50% when it was defined as the percentage of the hits to any of the 8 targets including KIT. The reason for the calculation limited to KIT was that the binding conformation of sunitinib was based on the complex with KIT (See “Preparation of compound conformation” in Methods). As a whole, screening by using AI-AAM on the basis of 5 references detected 11-75% of known compounds whose targets were the same as those of the reference compounds.

**Table 1.** The reference compounds and the hit compounds, with the extraction rate and the enrichment factor (EF) value of the hit compounds known to target the same protein as its reference.

| Reference compound |  | Hit compounds<br>by AI-AAM (n) |  | Reported<br>compounds<br>targeting the<br>same protein<br>(n) (A) | Extraction<br>rate (B/A)<br>(%) | EF value |
| --- | --- | --- | --- | --- | --- | --- |
| Name | Target | Same<br>target<br>(B) | Different<br>target |  |  |  |
| Aldosterone | NR3C2 | 7 | 37 | 11 | 63.6 | 86.5 |
| Testosterone | AR | 18 | 52 | 24 | 75.0 | 36.3 |
| Sildenafil | PDE5A | 2 | 11 | 3 | 66.7 | 33.3 |
| Sunitinib | KIT | 2 | 13 | 6 | 33.3 |  |
|  | Any target of sunitinib* | 10 | 5 | 20 | 50.0 | 8.3 |
| Celecoxib | PTGS2 | 2 | 0 | 18 | 11.1 | 95.2 |

\*CSF1R, FLT3, PDGFR, RET, FLT1, KDR, FLT4, KIT

Then, we calculated the enrichment factor (EF) (see “The enrichment factor” in Methods for more details). The random hit rates were approximately 0.01–0.25%, while the predicted hit rates were 1–8%. The predicted hit rates were apparently lower than the “extraction rates” shown in Table 1 because the denominators contained not only compounds that target the same proteins as references, but also those that target different proteins or have no target-molecule information. As shown in Table 1, the hit rate was improved by approximately 10–100 times (see Supplementary Fig. S1 online).

### Structure of the compounds identified using AI-AAM

The structural similarity of hits to the reference compounds was examined. In a graph representing AAM similarity on the vertical axis and Tanimoto coefficient on the horizontal axis, the hits were plotted (Fig. 3). For the compounds identified using aldosterone as a reference, Tanimoto coefficients were within the range of 0.1–0.6 (Fig. 3a). These values are rather low, which means that the hits include those with low structural similarity to the reference. Even for compounds whose known targets are the same as those of the reference compounds, Tanimoto coefficients are not always large. Moreover, most of the hits identified using sunitinib as a reference had a Tanimoto coefficient of less than 0.2 (Fig. 3b). In this way, the hits with low similarity could generally be identified, although there were some differences in the range of the Tanimoto coefficient depending on the reference compounds.

**Figure 3:**
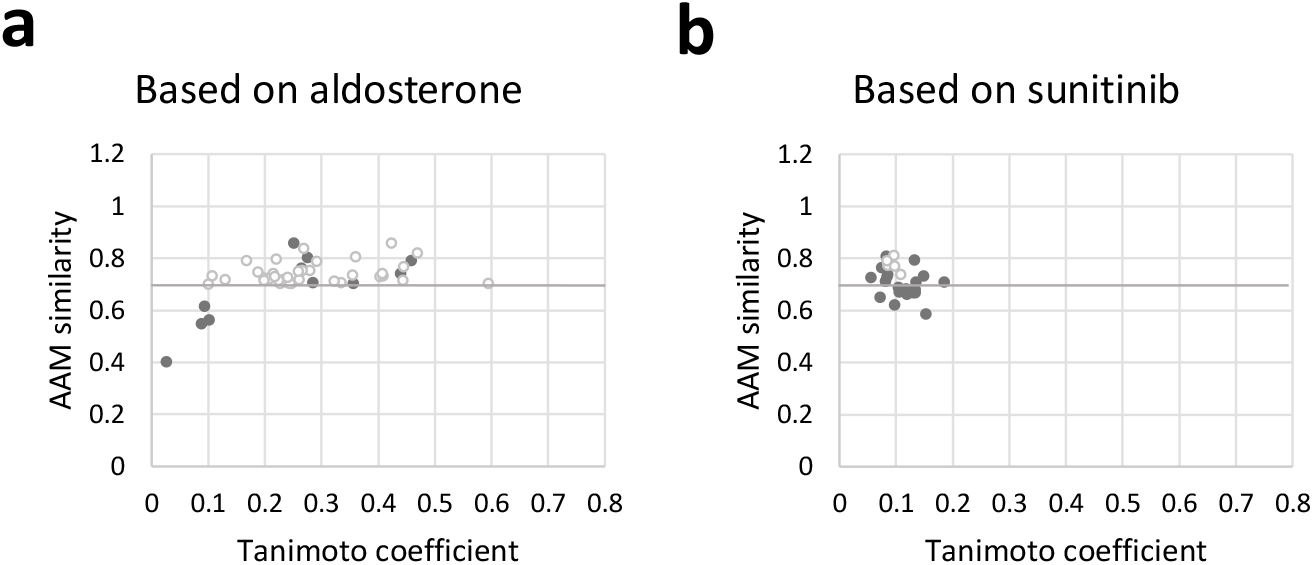
Tanimoto coefficients of hits and non-hits. Hits and non-hits based on aldosterone. (**a** target: NR3C2) or sunitinib (**b** target: KIT). The threshold value of AAM similarity as the boundary of hits and non-hits is 0.7. The hits include many compounds that target different proteins than the reference (white) as well as those which target the same proteins as the reference (black). Although the non-hits whose targets are the same protein as the references are also included (black), those known to target different proteins than the references are not shown.

### Specificity of screening using AI-AAM

As shown in Table 1, there were compounds whose known targets were the same as those of the reference compound among the hits, but more compounds known to target proteins different from the reference were identified as hits, suggesting that the latter also bind to the targets of their reference compounds (Fig. 4a). For example, among 44 hits identified with aldosterone as the reference, only 7 compounds were known to target NR3C2, the same protein targeted by the reference (Fig. 4b). In regards to the compounds screened on the basis of sunitinib, only 10 out of 15 hits were known to target any of the targets of sunitinib (Fig. 4b).

**Figure 4:**
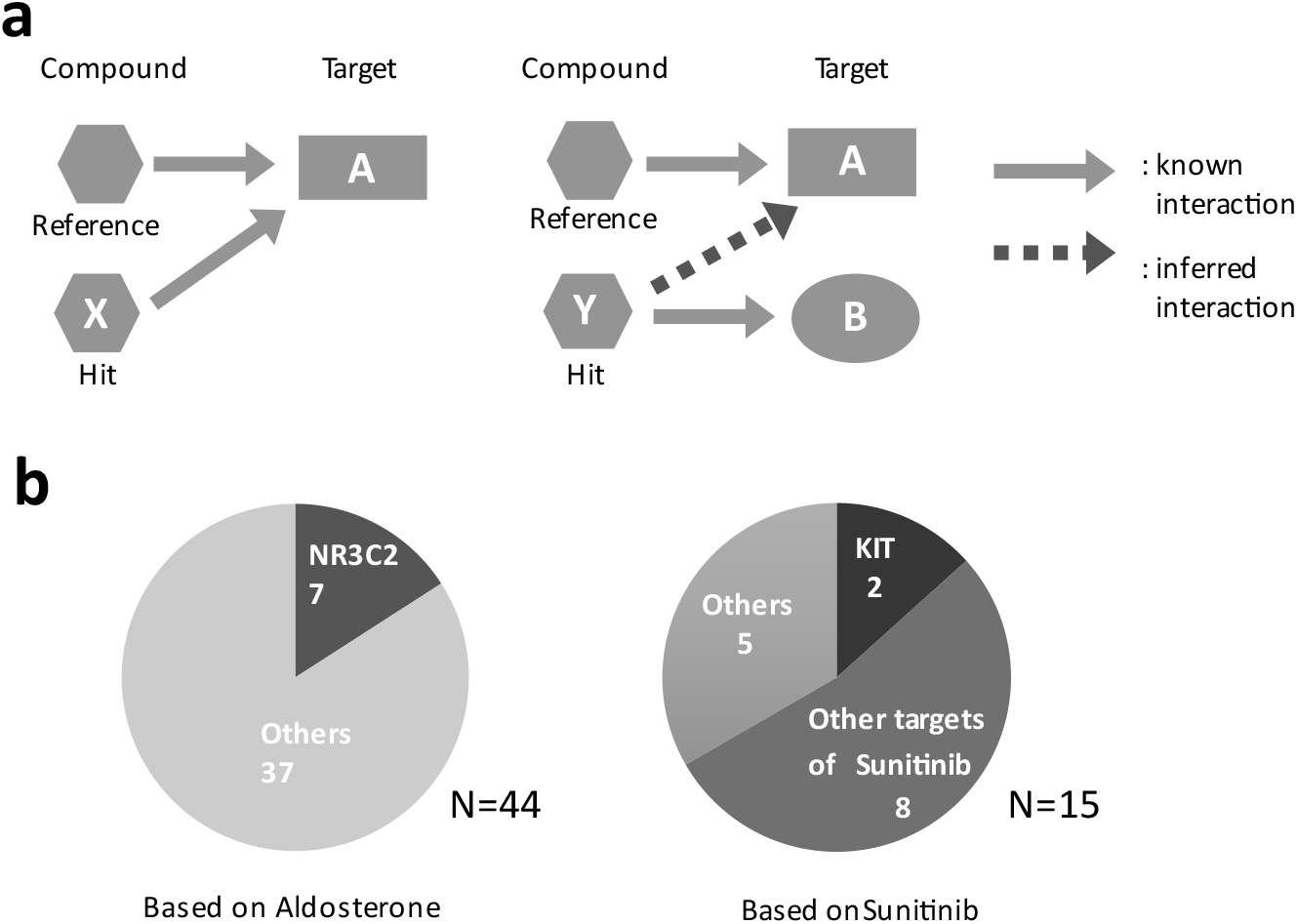
The targets of reference and hit compounds. **a** The reference and hit compounds and their target proteins. There are two types of hit-target interaction cases; cases where the known targets of the hit compounds are the same as those of the reference compounds and cases where the known targets of the reference and hit compounds differ. In the latter cases the targets of the reference compounds are presumed to be the unknown targets of hit compounds. **b** (Left) Hit compounds screened on the basis of aldosterone. Classification is based on the known targets: NR3C2 and others. (Right) Hit compounds screened on the basis of sunitinib. Classification is based on the known targets: KIT, other targets of sunitinib, and others.

An analysis to investigate the reason why the known targets of many hit compounds were different from those of the reference compounds was conducted. Among the compounds known to bind to the same targets as the reference compounds, some have the same biological function as the references, while others function in a different manner. We assumed that, as AI-AAM identified the compounds with the same function as the references, the comprehensiveness was underestimated. To verify this hypothesis, the functions of hits and non-hits were analyzed (Fig. 5, Supplementary Table S1 online).

**Figure 5:**
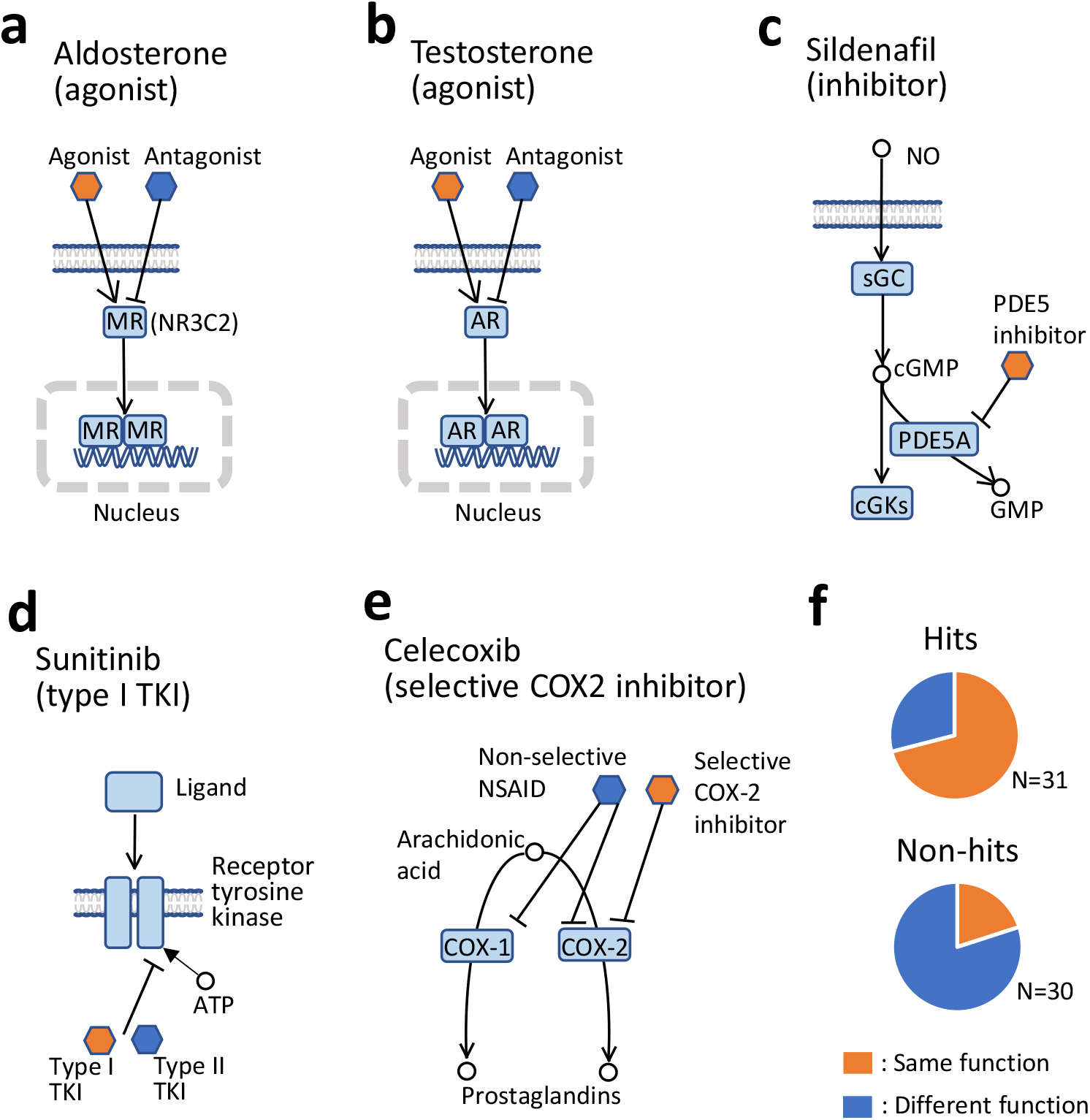
Pharmacological actions of reference compounds and the classification of hits and non-hits based on whether or not their function is the same as the reference. **a** Aldosterone targets NR3C2 as an agonist. NR3C2 is a ligand-activated nuclear transcription factor. NR3C2s in the inactive state reside primarily in the cytoplasm and are then transported to the nucleus, dimerize and form a transcription complex with DNA hormone response elements to initiate transcription of genes. The NR3C2 has affinity for both its primary physiological agonists and antagonists developed to treat diseases. **b** Testosterone targets the AR as an agonist. As with NR3C2, AR is nuclear transcription factor. Upon binding to endogenous ligands such as testosterone, AR translocates to the nucleus and regulate the transcription of genes. Apart from the endogenous ligands (agonists), exogenous ligands such as environmental chemicals and pharmaceuticals can also interact with AR as agonists or antagonists. **c** Sildenafi targets PDE5A. NO mediates its biological effects by activating sGC and increasing cGMP synthesis. As cGMP is degraded by PDE5A, its levels are maintained by inhibition of PDE5A by, for example, sildenafil. **d** Sunitinib is a type I tyrosine kinase inhibitor (TKI). Several kinases responsible for cell growth and proliferation are hyperactivated in various tumors. TKIs are the largest group of kinase-inhibiting small molecules. Most of the compounds, including sunitinib, act by blocking the ATP-binding site of the target molecule and the binding modes are classified as type I and type II depending on whether the compounds bind competitively with ATP using the ‘DFG-in’ (type I) conformation or the ‘DFG-out’ (type II) conformation. **e** Celecoxib targets COX-2 (gene symbol: PTGS2). NSAIDs (non-steroidal anti-inflammatory drugs) inhibit the enzyme cyclooxygenase (COX), which mediates the conversion of arachidonic acid to inflammatory prostaglandins. COX enzyme can exist in two forms: COX-1, the constitutive isoform; or COX-2, the inducible isoform. Selective COX-2 inhibitors are a subclass of NSAIDs that have a much greater affinity for the COX-2 enzyme, whereas non-selective NSAIDs inhibit both COX-1 and COX-2. **f** The ratios of the compounds with functions that are same as or different from reference compounds to hits or non-hits, which is based on the summary of the compounds identified using each of 4 compounds (i.e., aldosterone, testosterone, sunitinib and celecoxib).

As shown in Fig. 5a^19^, aldosterone is an agonist for nuclear receptor subfamily 3 group C member 2 (NR3C2, also known as MR). Eleven compounds are known to bind to NR3C2 in addition to aldosterone: 3 agonists and 8 antagonists. Seven hits were identified with aldosterone as the reference: 3 agonists and 4 antagonists. By using AI-AAM, agonists with the same function as the reference were obtained at a rate of 100%, while antagonists were identified at a rate of 50%.

Testosterone is an agonist for androgen receptor (AR) (Fig. 5b)^,20^. In addition to testosterone, 24 compounds are known to bind to AR: 14 agonists, 9 antagonists and a modulator. The number of hits identified applying AI-AAM based on testosterone was 18, 14 of which were agonists. The agonists, which have the same function as the reference, were obtained at a rate of 100%, while antagonists were identified at a rate of 44.4%. All 6 compounds that were not identified by using AI-AAM, even though these were known to target AR, were antagonists and a modulator. Chi-square test of independence shows a statistically significant relationship (p = 0.00162**) between the pharmacological action type (agonist or antagonist and the hit rate (see Supplementary Table S2 online).

Although phosphodiesterase 5A (PDE5A) inhibitors (Fig. 5c)^21^ identified using sildenafil as a reference were not classified according to their functions, their hit rate was originally high. Tyrosine kinase inhibitors (Fig. 5d)^22^ such as sunitinib could be classified into type I and type II^23, 24^. Although type I inhibitors, with the same function as sunitinib, were identified more often than type II, this was not statistically significant.

Non-steroidal anti-inflammatory drugs (NSAIDs), such as celecoxib, inhibit cyclooxygenase (COX) and are classified into selective COX-2 inhibitors and non-selective NSAIDs (Fig. 5e)^25, 26^. In addition to celecoxib, there are 7 compounds known to be selective for COX-2 (approved nomenclature for gene symbol: PTGS2). The number of non-selective NSAIDs was 10. Two hits identified using AI-AAM and celecoxib as the reference were selective inhibitors, and no non-selective NSAIDs were screened. Non-hit compounds included a cyclooxygenase-inhibiting nitric-oxide (NO) donator (CINOD) in addition to selective COX-2 inhibitors and non-selective NSAIDs. Moreover, there was a significant difference between the averages of AAM similarity of selective COX-2 inhibitors and non-selective NSAIDs regardless of hit or non-hit status (t-test, p = 0.0137*, see Supplementary Fig. S2 online).

The summary of hit and non-hit compounds identified using each of these 4 compounds (i.e., aldosterone, testosterone, sunitinib and celecoxib) as reference is shown in Table 2 and Fig. 5f. All compounds in this table target the same proteins as their reference compounds. Each hit and non-hit was further classified based on whether or not it had the same pharmacological mechanism of action as the reference. Chi-square test of independence showed a statistically significant relationship between two categorical variables: the result of screening (i.e., hits or non-hits) and function (p = 0.000065**). The screening using AI-AAM was highly selective of function.

**Table 2.** Contingency table for function data with row and column totals.

|  | Same function | Different function | SUM |
| --- | --- | --- | --- |
| Hits | 22 | 9 | 31 |
| Non-hits | 6 | 24 | 30 |
| SUM | 28 | 33 | 61 |

### Possibility of novel drug-target interactions suggested by AI-AAM

As mentioned above, there were compounds among the hits whose known targets were the same as those of the reference compound, while many compounds without information regarding whether they bind to the same protein as the references were also identified as hits (Table 1). We examined what kind of molecules were the targets when hits were known to target different proteins than the reference compound. As shown in Fig. 6, hits were classified on the basis of the biological functions of their known targets (i.e., nuclear receptor, enzyme, GPCR, and ion channel). The target information was obtained from DrugBank, and the target proteins were classified according to KEGG BRITE^27^.

**Figure 6:**
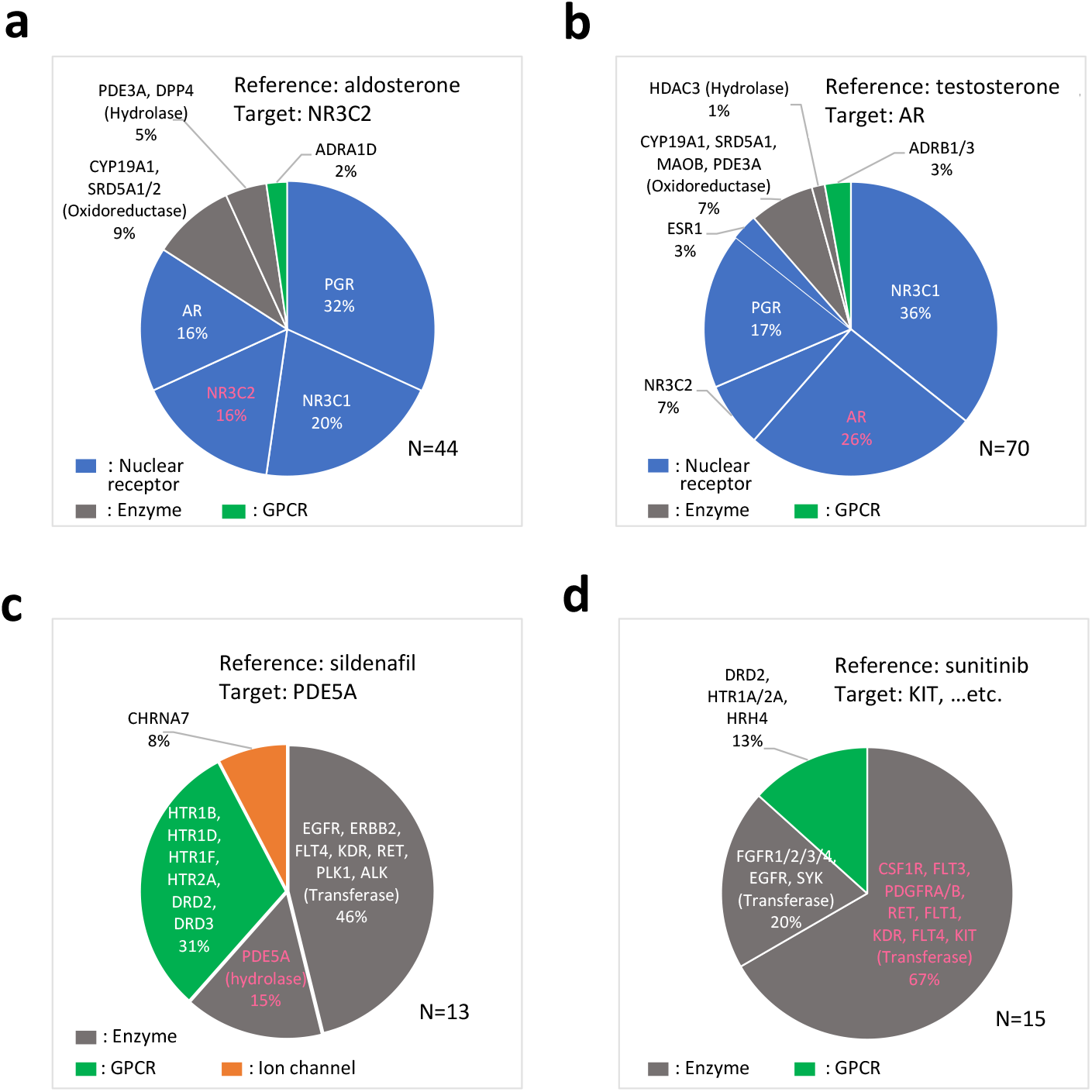
Hit compounds classified on the basis of the biological functions of their known targets Hit compounds screened on the basis of each reference compound. (**a** aldosterone, **b** testosterone, **c** sildenafil, **d** sunitinib). Classification is based on the biological functions of their known targets. Pink letters represent the same targets as the reference compounds.

The number of hits identified using AI-AAM with aldosterone as the reference was 44, with 37 known to target proteins other than NR3C2 (DrugBank, KEGG) (Fig. 6a). However, many of the known targets of the 37 compounds belonged to the same nuclear receptor family as NR3C2.

The number of hits screened with testosterone as the reference was 70, 52 of which were known to target different proteins other than AR (Fig. 6b). Most of these targets belonged to the same nuclear receptor family as AR, the targets of testosterone.

Sildenafil is known to target a hydrolase, PDE5A. Although 61% of the 13 hits obtained with sildenafil as the reference were known to target enzymes including hydrolases, the known targets of the remaining 31% and 8% of hits were GPCRs and ion channels, respectively (Fig. 6c). The known targets of sunitinib are some tyrosine kinases, and 67% of the 15 hits identified with sunitinib as reference were known to target the same tyrosine kinases as sunitinib. However, 20% and 13% of the hits were known to target other tyrosine kinases and GPCRs, respectively (Fig. 6d)

These results, together with those of other studies, indicate that many compounds identified through hits target multiple proteins of both the same and different families (see Discussion for more details).

## Discussion

In drug discovery, it is needed to obtain the improved compounds with excellent function and reduced side effects. To promote drug development for RIDs, we developed a new method of virtual screening, integrating AAM into scaffold-hopping, a LBVS technique. By applying this method to 5 compounds in DDrare, many hit compounds with diverse structures and the same affinity for a given target were obtained. As the EF values are equal to or greater than those of many SBVS techniques^28, 29, 30^, AI-AAM can be considered as the LBVS method to find the various compounds that target the same protein as the references with equal efficiency to SBVS methods. Moreover, our results show that XC608 identified with BIIB-057 as the reference has pharmacological activity equal to the reference. As well, for the compounds screened based on 5 compounds in DDrare, those with the same function as the reference tended to dominate the hits. This tendency was statistically significant regarding the compounds based on testosterone (target: AR) and celecoxib (target: PTGS2) (see Supplementary Table S2, Fig. S2 online). Singam et al.^31^ reported that agonists and antagonists of AR exhibit distinct binding modes: agonists form an H-bond with either Thr877 or Asn705, while this interaction is absent for antagonists. Other studies^32, 33, 34^ have reported that three amino acid differences between the COX-2 and COX-1 (gene symbol: PTGS1) active sites have major implications for the selectivity profile of inhibitors. In this way, pharmacological action is presumed to be closely related to the binding site of the target, especially some amino acids, and the differences in the function of the drugs are likely related to AAM similarity, as AI-AAM mainly considers the interaction between the candidate compound and each of the 20 amino acids. In other words, our results suggest that AI-AAM describes the interaction accurately, focusing on the essential qualities of interaction. In this way, if their known targets are the same as those of the reference compounds, the compounds likely are similar in function to the reference compounds, unlike those searched only on the basis of the target information in the databases. Moreover, even for such compounds, the structures are not always similar to the reference compounds, contributing to the expanded pharmacological space and the possibility of drug improvement by “hopping” from one scaffold to another^14^.

However, for the hits identified using 5 compounds as the reference, more compounds had known targets that differed from the targets of the reference compounds than those that targeted the same protein as the reference compounds. This suggests that the former also binds to the targets of reference compounds. Even if the known targets of the hits are different from those of the reference compounds, in many cases these belong to the same gene family. For example, all NR3C1 (GR), NR3C2, NR3C3 (PGR), and AR belong to the nuclear receptor family and have the same composition of functional domains, one of which, the steroid-binding domain (LBD), has sequence conservation to a certain degree^35^. Therefore, there may be some compounds active against multiple proteins of this family. However, the compounds that were inferred to interact with proteins of a different family were also not negligible. For example, the results showed that 2 of 15 hit compounds identified with sunitinib (target: receptor tyrosine kinases) as the reference were known to target aminergic GPCRs (i.e., DRD2, HTR2A, HTR1A, and HRH4); for 9 of 17 hits with BIIB-057 (target: SYK) as the reference, the known targets were GPCRs including aminergic GPCRs (i.e., HRH1, HTR1D, HTR1B, DRD2) (see Supplementary Fig. S3 online)^36^. This corresponds to Paolini’s report that a quarter of all of the compounds with multitarget activity (known as promiscuous compounds) are active across different gene families and aminergic GPCRs and protein kinases exhibit the greatest intra-as well as inter-gene family promiscuity^37^. Taken together, these results indicate the possibility of novel interaction between compounds and proteins, leading to multi-target information. The percentage of promiscuous compounds to the whole is reported to be approximately 20%^38, 39^. However, we believe that there are still many unknown interactions, as many ChEMBL data regarding the activity of compounds against targets are not yet reflected in the target information of databases such as DrugBank^40, 41^. The kinase assay in our study also showed that XC608 targets many kinases other than SYK, although the target of XC608, identified with BIIB-057 as the reference, was inferred to be merely SYK using AI-AAM, and it was validated by the experiment. It is probably another example to show the successful scaffold-hopping that the compounds with various sets of targets were screened on the basis of each reference compound.

In our previous study^42^, we invented a score *Rgene* for disease pairs sharing drug targets in RIDs, which represent a common mechanism of drug action underling drug repositionability. If a hit becomes known to share a target with the reference compound, the value of *Rgene* will rise. This means that the degree of drug repositionability between the indications of a hit and its reference are higher than that before the application of AI-AAM. The known target of a hit, prednisolone, screened with testosterone [target: AR] as reference is not AR, but NR3C1 (GR). However, one of the indications of prednisolone is muscular dystrophy, which is also known to be that of testosterone. Although AR is expressed at high levels in muscle^43, 44^, in the reports about corticosteroids for the treatment of Duchenne muscular dystrophy (DMD), this was not mentioned and the authors reported that the precise mechanism by which corticosteroids increase strength in DMD is not known^45, 46^. To confirm the possibility that prednisolone has pharmacological effects via the target of the reference compound, there is a need for further studies, but this may be an example of retrospective validation of this technique for DR. Recent approaches linked to network biology, the so-called ‘Network Pharmacology” are moving away from the current ‘one disease-one target-one drug paradigm of drug discovery’ that is becoming increasingly inefficient^47^. These approaches simultaneously target two or more proteins within disease-associated protein-networks^48, 49, 50^. The multi-target information that can be accumulated by our method will serve to clarify the mechanisms of action of drugs and, consequently, the disease mechanisms.

## Methods

### Reference compounds

#### For experimental validation of the technique

DDrare, a database of Drug Development for Rare diseases, was searched to identify proteins that are related to systemic lupus erythematosus (SLE), and SYK was finally selected as the target protein of the study. SLE is known as a RID, and there is a high need for new drugs to treat it. SYK is a non-receptor tyrosine kinase and was found to be related to many RIDs including SLE. Starting from a known SYK inhibitor candidate BIIB-057^51^, we aimed to discover other lead compounds.

#### For detailed analysis

DUD-E database, a database of 22,886 active compounds and their affinities against 102 targets^52^, was searched to find compounds that are also included in DDrare, and nine compounds were found. Four of them were removed because they were not covered by AI-AAM, such as a nucleic acid analogue (interaction with nucleic acids is important rather than with amino acids) and unsaturated fatty acid (too flexible). The remaining five compounds were used here as the reference compounds: aldosterone, testosterone, sildenafil, sunitinib and celecoxib,.

### Chemical library for virtual screenings

For both the prospective and the retrospective studies, we used a commercial library provided by NAMIKI SHOJI Co., Ltd. (https://www.namiki-s.co.jp/), which is composed of biologically active compounds including those in the clinical trial phase and approved drugs. All compounds in the library were preprocessed (desalted and normalized) by the MolStandardize module of RDKit v.2018.09.1, and 44,503 compounds preprocessed successfully were subjected to screening by AI-AAM.

### Calculation of AAM descriptors

AAM descriptors of each compound were calculated as distribution of centers of mass of amino-acid “probes” around the compound by using molecular dynamics (MD) simulation. However, its high computational cost makes it impractical for use in large-scale virtual screenings. Here we employed deep learning techniques to accelerate those calculations. The computational details are as follows:

#### Preparation of compound conformations

Experimentally determined binding conformations were used for calculating AAM descriptors of reference compounds except for BIIB-057. Cocrystal structures were downloaded from RCSB^53^ in PDB format under the PDB ID 2AA2 (aldosterone), 2AM9 (testosterone), 1UDT (sildenafil), 3G0E (sunitinib) and 3LN1 (celecoxib) and 3D structures of the ligands were extracted from the PDB files. Binding conformations of BIIB-057 and library compounds were unknown, and thus we considered 100 conformations generated by Discovery Studio 2020 (BIOVIA). Charge states at pH = 7.0 of all compounds including the reference compounds were also predicted by Discovery Studio.

#### Amino acid probes

In this study, not all 20 natural amino acids were considered but some were selected depending on the charge states of the reference compounds to reduce computational costs: asparagine, cysteine (deprotonated), phenylalanine, and threonine for monocationic reference compounds, and histidine (having protons both on the epsilon and delta nitrogen) was added for neutral reference compounds. To clearly highlight differences in AAM descriptors among amino acids, we removed their backbone chains and employed only side chains as the amino-acid probes.

#### Force fields

The generalized AMBER force field 2 (GAFF2)^54^ was employed to describe atomic interactions of reference and library compounds. Partial atomic charges were obtained by the restrained electrostatic potential (RESP) method^55^. The antechamber^56^ in the AmberTools19 package was used both to assign atom and bond types and to calculate the charges. For the RESP fit, electrostatic potentials of all the compounds were obtained from single-point quantum calculations at the HF/6-31G* level with the conformations optimized by the PM6 method. All of the quantum calculations were carried out by using Gaussian 16^57^. For amino acid probes, the ff14SB force field was employed. As a water model, TIP3P was used.

#### Calculations of AAM descriptors

In the beginning, we calculated AAM descriptors of various compounds as training data for deep learning by MD calculation. All MD calculations were performed using the gromacs-5.1.5 package^58^. As initial structures for the simulations, we considered pair structures of a compound and an amino acid probe which was placed randomly in the vicinity of the compound. A water box was created by tleap in AmberTools19 package where a minimum distance between any atom in the two molecules and an edge of a periodic box was set to 8.0 Å. A total of 100 pair structures were generated per compound, and all the structures were optimized and equilibrated with Berendsen thermostat and barostat. Here the coordinates of the compound were fixed, temperature T = 300 K, and pressure P = 1 atm. After a 100 ns production run with Nosé-Hoover thermostat and Parrinello-Rahman barostat, the AAM descriptor was calculated as an average distribution of a center of mass of the amino acid probe over 100 pair structures by using in-house software.

To learn AAM descriptors calculated by MD simulation, we adopted the pix2pix-type generative adversarial network model (https://doi.org/10.48550/arXiv.1611.07004). The original pix2pix was for 2D image data, and thus we extended it for 3D distribution data such as AAM descriptors. As explanatory variables, intermolecular potential energy surfaces (PES) between each probe atom and compounds were calculated. Note that PES calculations can be performed analytically and are therefore very fast. We confirmed that the obtained predictor can predict AAM descriptors of various compounds with sufficient accuracy. Here the number of training data (the number of compounds used for generating training data) was 100.

### Calculations of AAM similarity scores

AAM similarity scores were defined as the cosine similarities between AAM descriptors of the reference compounds and library compounds, and were calculated by the following process. We first defined a tensor of a reference compound for the AAM similarity score calculation as *g*_*a*_(*r*)θ(*r*), where *g*_*a*_(*r*) is an AAM descriptor of an amino acid probe *a* (*a* = 1 ∼ *N*_*a*_ is an index of amino acids, and *N*_*a*_ = 4 or 5) for a reference compound, *r* is a coordinate where the center of mass of the reference compound is taken as the origin, and

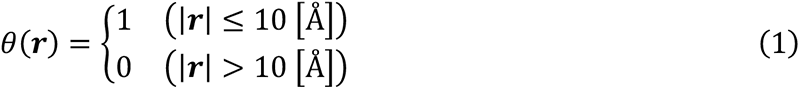

Now, the cosine similarity between AAM descriptors of the reference compound and a library compound can be defined as follows:

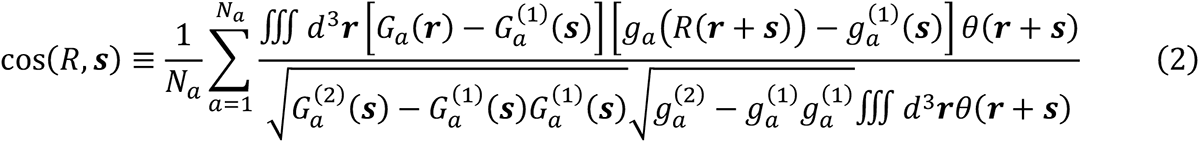

where *G*_*a*_(*r*) is an AAM descriptor of an amino acid *a* for the library compound, *s* = (*X, Y, Z*) is a translation vector, *R* = *R*(*α, β, γ*) is a rotation matrix and α∼γ are Euler angles, and

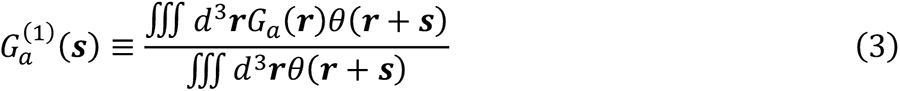

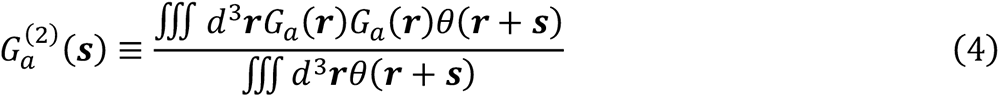

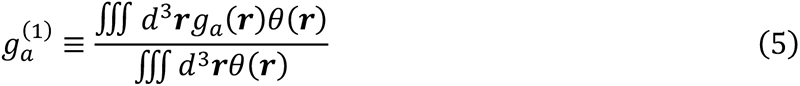

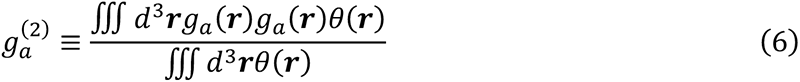

*s* and *R* were determined to satisfy the following condition by grid search with a step size of 1 Å for *X, Y*, and *Z* and 10 degrees for *α, β* and *γ*:

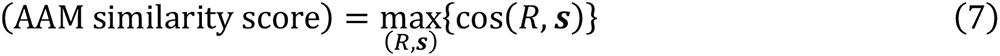

However, this requires high computational cost because the integration over *r* must be done *N*_*a*_ × *N*_*b*_ × *N*_*c*_ times (*N*_*R*_ and *N*_*s*_ are the numbers of grids for *R* and *S*). In a practical calculation, we reduced it by applying the singular value decomposition (SVD) method. Substituting

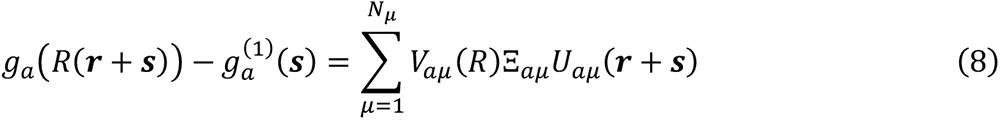

into eq.(2), the following expressions can be obtained:

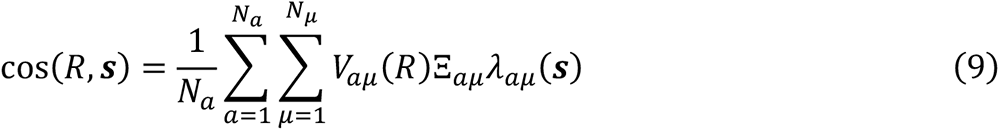

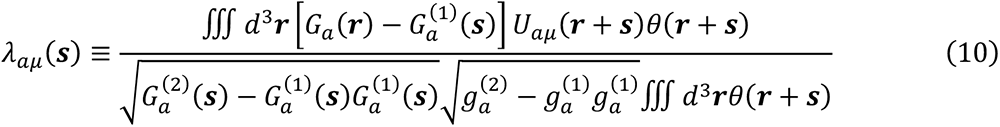

where a cutoff *N*_μ_ was determined so that a summation of explained variance ratios exceeded 0.99. It is clear from these expressions that the number of integrations over ***r*** was decreased from *N*_*a*_ × *N*_*R*_ × *N*_*s*_ to *N*_*a*_ × *N*_*μ*_ × *N*_*s*_.

We calculated AAM similarity scores of all pairs of the reference compounds and library compounds, and extracted compounds whose AAM similarity scores were > 0.7 as candidate compounds.

### Calculations of Tanimoto coefficients

Morgan fingerprint with radius 2 and 2048 bits was used for calculations of Tanimoto coefficients between library and reference compounds. All the calculations were done by using RDKit v.2018.09.1.

### The enrichment factor (EF)

EF values are commonly used in virtual screening evaluation as accuracy metrics. The EF value is defined as the ratio between the predicted hit rate and the random hit rate^28, 29^. At first, the hit rate of the compounds known to target the same proteins as their reference was estimated when AI-AAM was not applied. To calculate the hit rate, the number of compounds that were known to target the same proteins as the reference and were contained in the subset of the NAMIKI library, which consists of the compounds having the same electrical charge as the reference, was counted. This was then divided by the total number of the compounds contained in the library (“random hit rate”). Subsequently, the hit rate of the compounds whose known targets were the same as those of the reference compound was estimated when AI-AAM was applied. The value was the number of the hit compounds known to target the same proteins as the reference divided by the total number of hits (“predicted hit rate”). EF values were calculated by dividing the “predicted hit rate” by the “random hit rate” for each reference compound.

### In vitro SYK kinase assay

The experiments to evaluate the inhibitory activity levels of compounds were carried out according to the manufacturer’s instructions on the SYK Kinase Enzyme System (Promega, Wisconsin, USA) and ADP-Glo^TM^ Kinase Assay (Promega). By measuring luciferase activity using the Ensight Multimode Plate Reader (PerkinElmer), the inhibition rate was calculated by comparing the OD value to the negative and positive control wells.

### Mobility shift assay

Kinase profiling of BIIB-057 and XC608 against a panel of 24 kinases was performed to examine the selectivity by using mobility shift assay at Carna Biosciences, Inc. (Kobe, Japan) (https://www.carnabio.com/english/product/msa.html). Drug concentrations were both set to 50 nM, and ATP concentrations were approximately equal to the Km value for each kinase.

### Statistical analysis

All statistical analyses were performed with Excel. To identify the relationship between two categorical variables, Chi-square test was performed. To calculate the probability of significant difference between two groups, two-tailed t-test (unpaired) was performed. *P < 0.05 were considered as statistically significant. Before the t-test, F-test was performed to determine if the two samples had equal variance.

## Data Availability

The datasets generated and analyzed during the current study are available from the corresponding author on reasonable request.

## Acknowledgements

We thank Yuko Usami for supporting the original data construction. This study was supported in part by a grant from the Japan Society for the Promotion of Science (JSPS, Grant Number 20K12056).

## Author contributions

Conceptualization: R.S. and T.K.; Design: R.S. and T.K.; Acquisition and construction of data: R.S., M.T., J.N., K.T., S.O., and K.I.; Analysis of data: R.S., M.T., T.K., J.N., K.T., S.O., and K.I.; Interpretation of data: R.S. and T.K.; Supervision: T.K.; Writing manuscript: R.S., M.T., and T.K. All authors read and approved the final manuscript.

