## Supplementary File for "A novel *in silico* scaffold-hopping method for drug repositioning in rare and intractable diseases"

Supplementary Table S1: The classification of hits and non-hits based on whether or not their pharmacological action type is same as the reference.

| **Reference compound** | | | **The compounds that target the same protein as the reference compound** | | |
|---|---|---|---|---|---|
| Name | Target | Pharmacological action type | Pharmacological action type | Hits (N) | Non-hits (N) |
| Aldosterone | NR3C2 | **Agonist** | **Agonist** | 3 | 0 |
| | | | Antagonist | 4 | 4 |
| Testosterone | AR | **Agonist** | **Agonist** | 14 | 0 |
| | | | Antagonist/ Modulator | 4 | 6* |
| Sildenafil | PDE5A | Inhibitor | Inhibitor | 2 | 1 |
| Sunitinib | KIT | **Type I inhibitor** ** | **Type I inhibitor** | 3 | 1 |
| | | | Type II inhibitor | 1 | 3 |
| | | | Unknown | 6 | 6 |
| Celecoxib | PTGS2 | **Selective COX-2 inhibitor** | **Selective COX-2 inhibitor** | 2 | 5 |
| | | | Non-selective NSAIDs | 0 | 10 |
| | | | CINOD | 0 | 1 |

*Antagonist: 5, Modulator: 1

**Wang, B et al. (2021)[23], Zhao, Z et al. (2014)[24]

Supplementary Table S2: Contingency table for function data of hits and non-hits screened on the basis of testosterone.

| | Agonist | Antagonist | SUM |
|---|---|---|---|
| Hits | 14 | 4 | 18 |
| Non-hits | 0 | 5 | 5 |
| SUM | 14 | 9 | 23 |

**Supplementary Figure S1: The hit rate of the compounds targeting the same protein as the reference compound when AI-AAM is not applied or applied.**

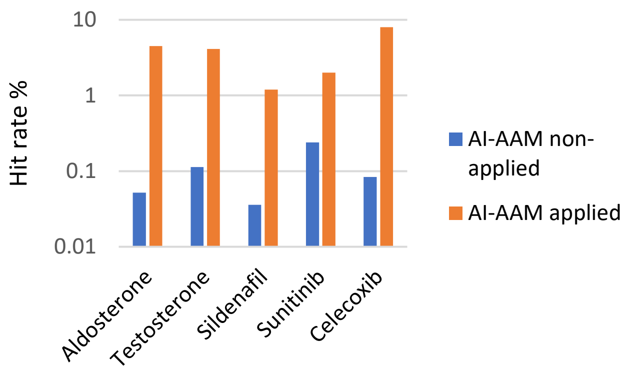


The graph uses logarithmic scale on the vertical axis.

**Supplementary Figure S2: AAM similarity of selective COX2 inhibitors and non-selective NSAIDs identified with celecoxib as a reference.**

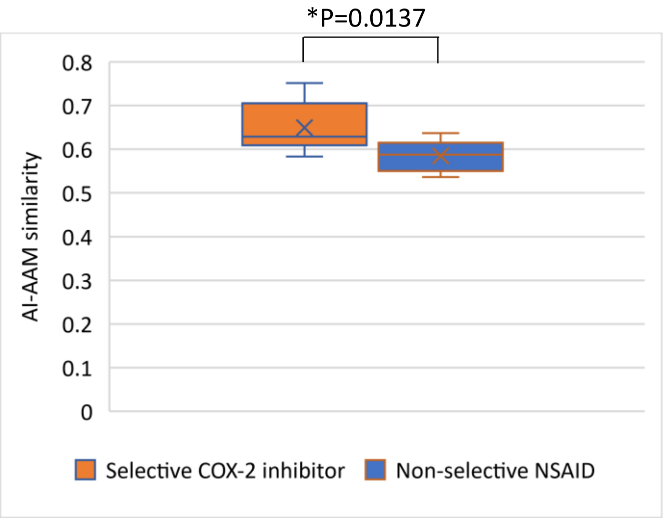


Each box-and-whisker plot shows the five-number summary of a set of data for AI-AAM similarity of selective COX-2 inhibitors (n=7) and non-selective NSAIDs (n=10), respectively. A box is drawn from the first quartile to the third quartile. A vertical line goes through the box at the median. The whiskers go from the ends of the box to the minimum or maximum values. P value was calculated by two-tailed unpaired t-test. The asterisk indicates the statistically significant difference ($p < 0.05$).

**Supplementary Figure S3: Hit compounds (reference: BIIB-057) classified on the basis of the biological functions of their known targets**

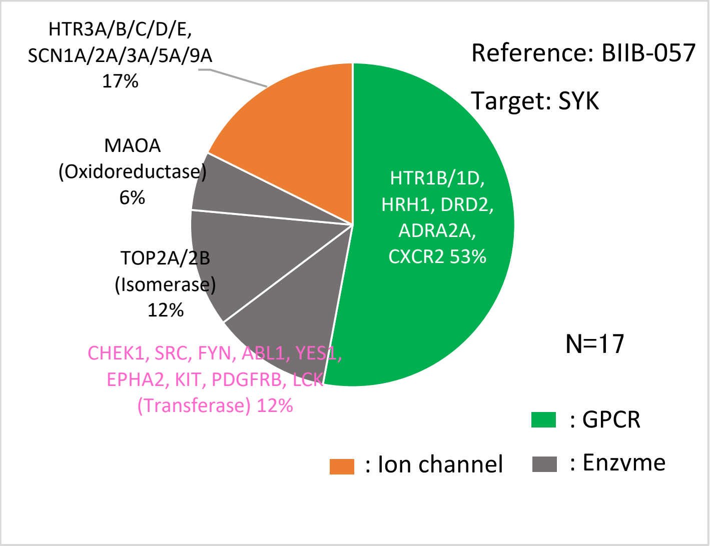


Pink letters represent the same targets as the reference compounds.
